# Cerebello-Thalamic Spike Transfer via Temporal Coding and Cortical Adaptation

**DOI:** 10.1101/2020.01.19.911610

**Authors:** Carmen B. Schäfer, Zhenyu Gao, Chris I. De Zeeuw, Freek E. Hoebeek

**Author notes:** Correspondence: Prof. Dr. Freek E. Hoebeek, Department for Developmental Origins of Disease, Wilhelmina Children’s Hospital, room KC.03.071.0, Lundlaan 6, 3584 EA, Utrecht, Netherlands.

## Abstract

Orchestrating the ensemble of muscle contractions necessary for coordinated movements requires the interaction of cerebellar, thalamic and cerebral structures, but the mechanisms underlying the integration of information remain largely unknown. Here we investigated how excitatory inputs from cerebellar nuclei (CN) and primary motor cortex layer VI (M1 L6) neurons may regulate together the activity of neurons in the mouse ventrolateral (VL) thalamus. Using dual-optogenetic stimulation and whole-cell recordings *in vitro* we were able to specifically activate the CN and M1 pathways and study their differential impact. We found that VL spiking probability is effectively determined by a pause in CN stimuli, whereas VL membrane potential can be modulated subthreshold by M1 L6 input. Upon mild depolarization of the VL membrane potential, repetitive CN stimulation evokes at best single action potential firing, whereas more negative membrane potentials increase VL spiking probability. Moreover, whereas high-frequency cerebellar activity attenuates thalamic spiking, pauses in cerebellar activity re-activate thalamic spiking. In contrast, facilitating inputs from cerebral cortex modulate thalamo-cortical spike transfer via fluctuations in the membrane potential. The fine-tuning by cerebellar and cerebral activity allows the motor thalamus to operate as a low-pass filter for cerebellar activity, generating sparse but precisely timed outputs for the cerebral cortex.

## Introduction

Successful motor learning requires an estimation of the sensory consequences of a motor plan and the integration of error signals into ongoing sensorimotor processing (Ramnani 2006; Brooks et al. 2015). This complex task requires the communication of multiple brain areas, such as the cerebellum, thalamus and motor cortex. The execution of acquired movements is mediated by cerebellar computation, in that genetic and functional lesions throughout various cortical and deeper cerebellar regions are known to disrupt the execution of motor behavior (De Zeeuw and Ten Brinke 2015). Subsequently, the cerebellar output, embodied by projection neurons in the cerebellar nuclei (CN), is integrated in the ventrolateral nucleus (VL) of the thalamus and from there relayed to various layers of the motor cortex (Teune et al. 2000; Kuramoto et al. 2009; Proville et al. 2014; Svoboda and Li 2017; Gornati et al. 2018). In addition to cerebellar information, VL neurons receive excitatory input from cortical layer 6 neurons of the primary motor cortex (M1 L6) (Yamawaki and Shepherd 2015; Jeong et al. 2016). The interaction of subcortical and cortical inputs has been shown to determine thalamic output (Mease et al. 2014) and is therefore of key importance to improve our understanding on cerebello-thalamic information processing in motor control.

At rest, the baseline firing rates of CN range between 30 and 100 Hz (Hoebeek et al. 2010), while VL and cortical L6 neurons fire at low frequencies between 5 and 20 Hz (Lamarre et al. 1971; Vitek et al. 1994; Beloozerova et al. 2003; Marlinski et al. 2012; Olsen et al. 2012; Proville et al. 2014). Once movement execution starts, CN neurons evolve into phasic patterns including high-frequency bursts of spiking (Antziferova et al., 1980; Armstrong and Edgley, 1984; Ohmae et al., 2013; Sarnaik and Raman, 2018). The well-timed and rapid alterations in CN activity patterns are destined to adapt thalamo-cortical spiking in VL (Proville et al., 2014). It remains an open question how CN-VL synaptic transmission, which is subject to paired-pulse depression and L6-VL transmission (Gornati et al. 2018), interact in individual neurons and corroborate the VL spiking patterns that are characterized by burst-pause and tonic spiking. In more detail, it is unclear how the pauses in the CN spiking, which are thought to decode the timing of specific sensory events (De Zeeuw et al. 2011), can affect VL neuronal activity and how cortico-thalamic modulation of VL membrane potential affects these supposed responses.

Answering these open questions regarding cerebello-cerebral interactions at the network level requires a detailed understanding of the cerebello-thalamic and cortico-thalamic inputs at the cellular level. In the current study we investigated the synaptic interplay at VL neurons that receive both CN and M1 L6 inputs in an *in vitro* preparation, which allows us to pharmacologically and optogenetically control the network activity. While blocking inhibitory synaptic transmission we recorded VL membrane potentials in whole cell patch clamp mode in combination with dual-optogenetic stimulation techniques to selectively stimulate CN and M1 L6 axons. Our results show that pauses in CN spiking are a determinant of VL output and that M1 L6 inputs potently modulate the cerebellar induced spiking in VL neurons, together constructing a low-pass filter that can be fine-tuned in a timing-dependent manner.

## Material and Methods

### CONTACT FOR REAGENT AND RESOURCE SHARING

Information and requests for resources and reagents should be directed to and will be fulfilled by the lead contact, Freek E. Hoebeek.

#### Experimental Model Details

All experiments were performed in accordance with the European Communities Council Directive. All animal protocols were approved by the Dutch national experimental animal committee (DEC). For all experiments Tg^(*Ntsr1-cre*)GN220Gsat^ (*Ntsr1-Cre*) transgenic mice were used (Gong et al. 2007). The colony was originally purchased from the MMRRC repository and maintained by backcrossing with C57Bl/6^OlaHsd^ mice. The genotype was tested by PCR reaction using toe-tissue gathered at postnatal (P) 7-10. For physiology experiments *Ntsr1-Cre* mice were used from 21 days of age and for anatomical experiments from 60-120 days of age.

#### Stereotaxic Injections and Viral Vectors

For surgery mice were anaesthetized with isoflurane (5% in 0.5 L/min O_2_ during the induction and 1.5% in 0.5L/min O_2_ for maintainance). After skin incision, local anaesthesia was maintained using lidocaine (10%, local application) and buprenorphine (50 µg/kg bodyweight) was applied for analgesia. Craniotomies of 0.5-2 mm were established above the injection sides. For injections to M1, approximately 200nl of adeno-associated virus (AAV) was injected to each of the following stereotaxic coordinates relative to bregma and midline (x, y; in mm): (1) 1.5, 1; (2) 1.5, 1.25; (3) 1.5, 1.5 at −0.9 depth from the dura. For injections to the CN, 200 nL of AAV was injected 2 mm posterior to lambda, 2 mm lateral to the midline at a depth of −2 mm from the dura and on the contralateral side to M1 injections. For optogenetic stimulation experiments, AAV2.9-hSyn-FLEX-ChrimsonR-tdT (provided by Prof. Bryan Roth through the UNC vector core) was injected to M1 and AAV2.9-hsyn-ChR2(H134R)-EYFP was delivered to CN (provided by Prof. K. Deisseroth through the UNC vector core). For all optogenetic stimulation experiments, both injection sides were placed unilaterally with the cerebellar injection side contralaterally to the motor cortical injection side. Injection spots of dually injected animals are represented in **Supplementary Fig.1**. Mice that showed ChR2 expression in their vestibular nuclei neurons were excluded from analysis. For anterograde tracing experiments we used AAV constructs (chimeric serotype 1 and 2) carrying CAG_Synaptophysin_eGFP as well as CAG_Synaptophysin_mOrange (kindly provided by Prof. T. Kuner, Heidelberg University) and for retrograde tracing we injected 1% Cholera toxin subunit B (CTB).

#### Preparation of Acute Slices

Following 3-6 weeks of incubation isoflurane-anesthetized mice were decapitated, their brains were quickly removed and placed into ice-cold slicing medium containing (in mM): 93 NMDG, 93 HCl, 2.5 KCl, 1.2 NaHPO_4_, 30 NaHCO_3_, 25 Glucose, 20 HEPES, 5 Na-ascorbate, 3 Na-pyruvate, 2 Thiourea, 10 MgSO_4_, 0.5 CaCl_2_, 5 N-acetyl-L-Cysteine (osmolarity 310 ± 5; bubbled with 95% O_2_ / 5% CO_2_). Next, 250 μm thick coronal slices were cut using a Leica vibratome (VT1000S). For the recovery, brain slices were incubated for 5 min in slicing medium at 34±1°C and subsequently for ∼40 min in artifical cerebrospinal fluid (ACSF; containing in mM: 124 NaCl, 2.5 KCl, 1.25 Na_2_HPO_4_, 2 MgSO_4_, 2 CaCl_2_, 26 NaHCO_3_, and 20 D–glucose, osmolarity 310±5; bubbled with 95% O_2_ / 5% CO_2_) at 34±1°C. After recovery brain slices were stored at room temperature before the experiments started. For the confirmation of the injection spot, motor cortices and hindbrain were conserved in 4% paraformaldehyde (PFA).

#### Electrophysiology and Photostimulation

For all recordings, slices were bathed in 34±1°C ACSF (bubbled with 95% O_2_ / 5% CO_2_) and supplemented with 100 µM picrotoxin to block for GABAergic inputs, e.g., from the reticular nucleus in the thalamus. Whole-cell patch-clamp recordings were performed using an EPC-10 amplifier (HEKA Electronics, Lambrecht, Germany) for 20-60 min and digitized at 50 kHz. Recordings were excluded if series or input resistances (RS and RI, respectively) varied by >25% over the course of the experiment or if RS exceeded a maximum of 25 MΩ. Voltage and current clamp recordings were performed using borosilicate glass pipettes with a resistance of 3-6 MΩ when filled with K^+^-based internal (in mM: 124 K-Gluconate, 9 KCl, 10 KOH, 4 NaCl, 10 HEPES, 28.5 Sucrose, 4 Na_2_ATP, 0.4 Na_3_GTP (pH 7.25-7.35; osmolarity 295 ± 5)). All recording pipettes were supplemented with 1 mg/mL Biocytin (Sigma-Aldrich, St. Louis, USA) to allow histological staining (see below). When necessary internal solution was supplemented with QX-314 (10 mM, Sigma-Aldrich, St. Louis, USA), a blocker of voltage-gated Na^+^-channels, to prevent escape spikes in response to photocurrents in M1 and CN neurons. Current clamp recordings were corrected offline for the calculated liquid junction potential of −10.3 mV.

Dual optogenetic stimulation was induced using a pE-2 (CoolLED, Andover, UK) with LED wavelengths at 470 nm and 585 nm in combination with a dichroic mirror at 580 nm (filterset 15 without excitation filter, Carl Zeiss, Jena, Germany) and a 40X objective (Carl Zeiss). Light intensities were recorded by collecting photons across an area of 1 cm^2^ (PM400 Optical Power Meter, Thorlabs, Newton, USA) and the power was back calculated to the area of the focal spot to determine stimulation intensities. The photon flux was calculated by converting the irradiance via the following formula:

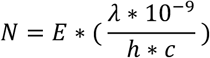

in which N is the number of photons, E is the irradiation in [W/m^2^], λ is the wavelength in [nm], h is the Max-Planck constant and c is the speed of light. Full-field dual optogenetic stimulation with 585 nm was applied for 15 ms with an intensity of 1.66 mW/mm^2^ and a photonflux of ∼4.8*10^24^ photons/ms/m^2^, while stimulations at 470 nm were applied for 1ms with intensities ranging from 0.99–7.65 mW/mm^2^ (maximally ∼18.08*10^24^ photons/ms/m^2^). The photostimulation resulted in maximally inducible response amplitudes from CN and M1-L6 fibers. To ensure that we recorded AP-driven neurotransmitter release a portion of the experiments were concluded by bath application of 10 μM tetrodotoxin (TTX, Tocris, Bristol, UK).

#### Histology

For the histological reconstruction of the patched neurons, the brain slices were placed in 4% paraformaldehyde (PFA; in 0.1 M PB and pH 7.3) for 3-5 days. After rinsing the slices with 0.1 PB, they were placed in 10 mM Na-citrate at 80°C for 3 h and afterwards blocked for 2 h at RT (10% normal horse serum (NHS) and 0.5% Triton-X100 in PBS). Synaptic varicosities were labeled with primary antibody for vesicular glutamate transporter 1 (vGluT1; 1:2000, 8 days incubation, Guinea Pig, MilliPore Biosciece Research Reagent, Kenilworth, USA) and vesicular glutamate transporter 2 (vGluT2; 1:2000, Guinea Pig, MilliPore Bioscience Research Reagent, 2 days incubation) in 2% NHS and 0.5% Triton-X100. As secondary antibody we used anti Guinea Pig A405 (1:400, Jackson Immunoresearch, Bar Harbor, USA) while biocytin-filled neurons were visualized by overnight incubation with Streptavidin-Cy5 conjugated antibody (1:400, Jackson Immunoresearch).

#### Image Acquisition and Analysis

Widefield images and confocal images were acquired on a LSM 700 microscope (Carl Zeiss) by using a 20X/0.30 NA and 63X/1.4NA objective, respectively. For the morphological reconstruction of vGluT1/2 staining, ChR2 fibers, ChrimsonR fibers and biocytine-filled cells the following excitation wavelengths and fluorophores were used (in order): 405 nm (Alexa 405), 488 nm (YFP), 555nm (tdTomato) and 639 nm (Cy5). High-resolution images were oversampled with 30 nm pixel and 130 nm z-step size and subsequently deconvoluted by using Huygens Software (Scientific Volume Imaging, Hilversum, The Netherlands).

#### Quantification and Statistical Analysis

All sweeps were recorded 3-10 times and averaged. Data analysis was performed using Clampfit software (HEKA Electronics) or custom written routines in IGOR Pro 6.21 (Wavemetrics, Lake Oswego, Oregon, USA). For trains of stimuli, the peak amplitude of each evoked postsynaptic current/potential (EPSC/EPSP) was detected relative to baseline. Normalized EPSC amplitudes within the train were divided by the amplitude of the first evoked response. The steady state amplitude of higher frequency responses as well as the spike probability was calculated by averaging the response during the last 100 ms and 500 ms within the train, respectively. The CV is defined by the ratio of the standard deviation to the mean. For spike recordings in response to CN-stimulation, cells with a CV>0.2 were excluded from the analysis. All data was tested for normal distribution with the Kolmogorov-Smirnov test. For statistical comparisons Mann-Whitney-test, Wilcoxon matched-pairs signed rank test or Friedman-test with correction for missing values by pairwise exclusion were applied instead. For trend analysis the Cochran Armitrage test was applied. For statistical analyses GraphPad PRISM, SPSS and R software packages were used. All datasets were corrected for multiple comparisons and are represented as mean ± standard error of the mean (SEM).

#### Data and Software availability

Data and software codes will be made available upon consent of the lead author. Please contact for requests.

## Results

To identify the cell-physiological mechanisms that enable VL neurons to integrate CN with M1-L6 inputs we first evaluated the effect of CN stimulation on the spike output of VL neurons in acute slices (**Figure 1A**, injection sites depicted in **Supplementary Figure 1**). Upon stimulation of ChR2-expressing CN fibers with a single 1 ms light pulse (470 nm) we recorded an EPSC of 1107±143 pA carrying a charge of 4745±543 pA*ms (**Supplementary Figure 2A-F**; for statistical analyses see **Supplementary Table**). Stimulating CN fibers at 20, 35 and 50 Hz resulted in a short-term decrease of EPSC amplitudes (**Supplementary Figure 2G-J**). When evaluating the spiking at average resting potentials of −73.4±0.4 mV, VL neurons responded with an initial burst of action potentials followed by a steady state of 0.20±0.08 and 0.15±0.07 following the 20 and 50 Hz stimulus trains, respectively (**Figure 1B,C**; *P*=0.0625, Wilcoxon-signed rank (WCR) test, n=25; **Supplementary Figure 2K-M**). Next, we tested the thalamic response to brief pauses in the CN stimulation, which commonly occur during motor behaviors (Antziferova et al. 1980; Armstrong and Edgley 1984; Ohmae et al. 2013; Sarnaik and Raman 2018). Interestingly, increasing the pause length recovered the attenuation of the CN-evoked VL responses and increased the spiking number (**Figure 1D**; 1.39±0.46 and 2.43±0.65, respectively; Friedman-test, 100 ms vs 400 ms: *P*=0.019). The recovery of the EPSC was modulated by increasing pause lengths, but not by stimulation frequency (**Supplementary Figure 1G, O, P**). These data suggest that VL neurons function as a low-pass filter for the high-frequency CN inputs, with unique properties to translate pauses in CN spiking into well-timed thalamic outputs.

**Figure 1.**
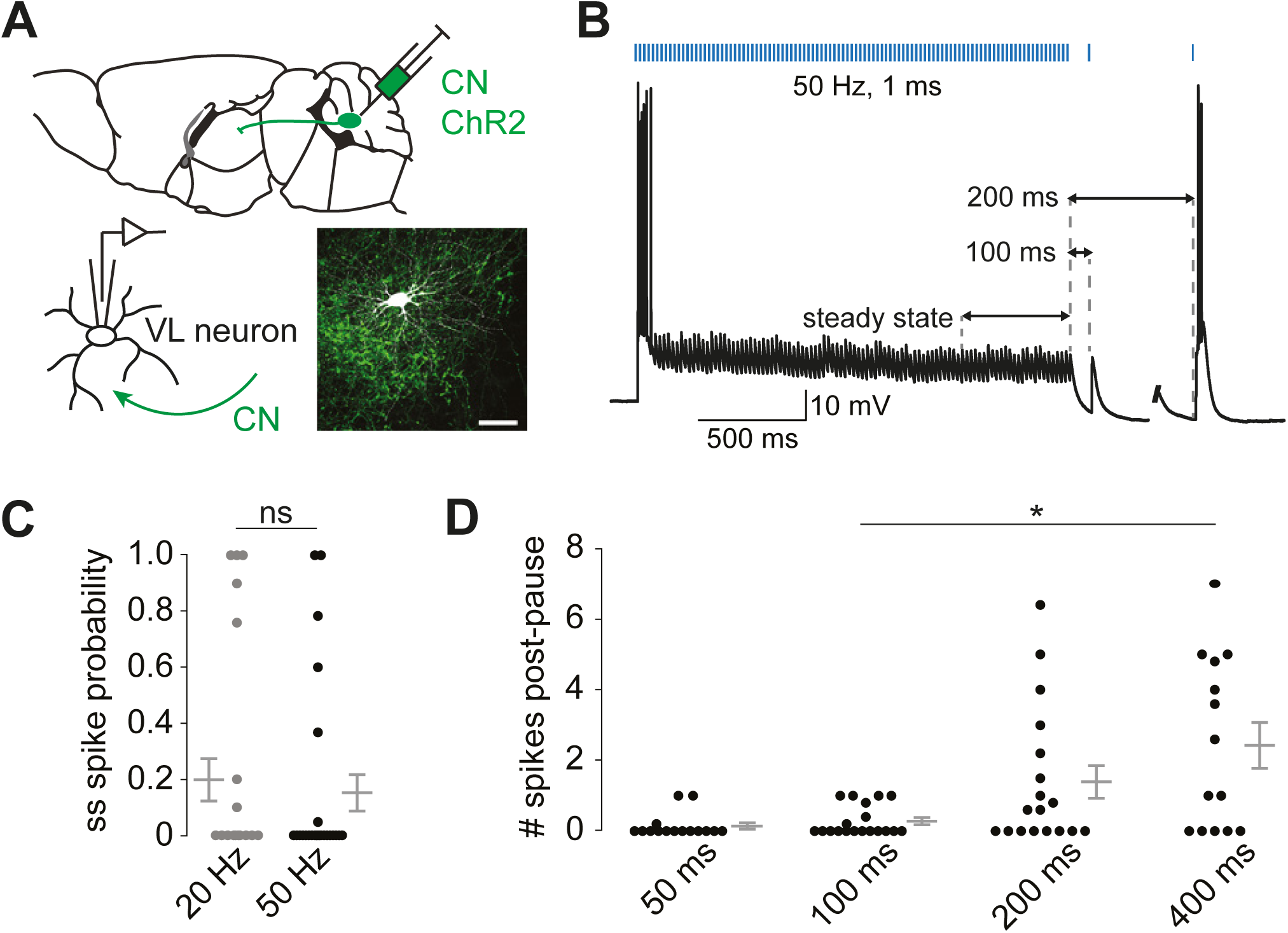
Sensitivity of VL thalamic spike responses to pauses in CN stimulation. (A) Illustration of a patch-clamped neuron embedded in ChR2-expressing CN fibers (green, scale bar: 50 µm). (B) Example trace of VL responses to 50 Hz stimulation followed by 100 or 200 ms pauses at resting potential. (C) Cerebellar-evoked EPSPs fail to trigger thalamic spiking during steady state (ss), which recovers with increasing pause length (D). ‘ns’ indicates not significant and * P<0.05.

To establish the convergence of CN and M1-L6 inputs on VL neurons that in turn project to motor cortex we injected adeno-associated virus (AAV) expressing synaptophysin-mCherry into CN and a mix of AAV expressing synaptophysin-GFP and cholera toxin-B (CTB, retrograde tracer) into M1. Synapses originating from CN and M1-L6 neurons converged within close proximity on CTB labelled VL neurons, which innervate M1 (**Figure 2A-C**). This connectivity pattern suggests that cerebellar output is integrated in this topographically organized cortico-thalamo-cortical loop between M1 and VL. Likewise, morphological convergence of ChrimsonR-tdtomato positive fibers from motor cortical layer 6 and Channelrhodopsin 2 (ChR2)-YFP positive fibers from CN in cerebellar-recipient VL suggests the convergence of dual inputs from M1-L6 and CN on the same cell and dendrite (**Figure 2D,E**). Moreover, co-labelling the M1-L6 and CN axons with vGluT1 and vGluT2 shows their cortical and subcortical origin, respectively (**Figure 2D,E**). These data indicate that VL receives both subcortical signals from CN and cortical feedback from motor cortical layer 6, generating opportunities for heterosynaptic integration.

**Figure 2.**
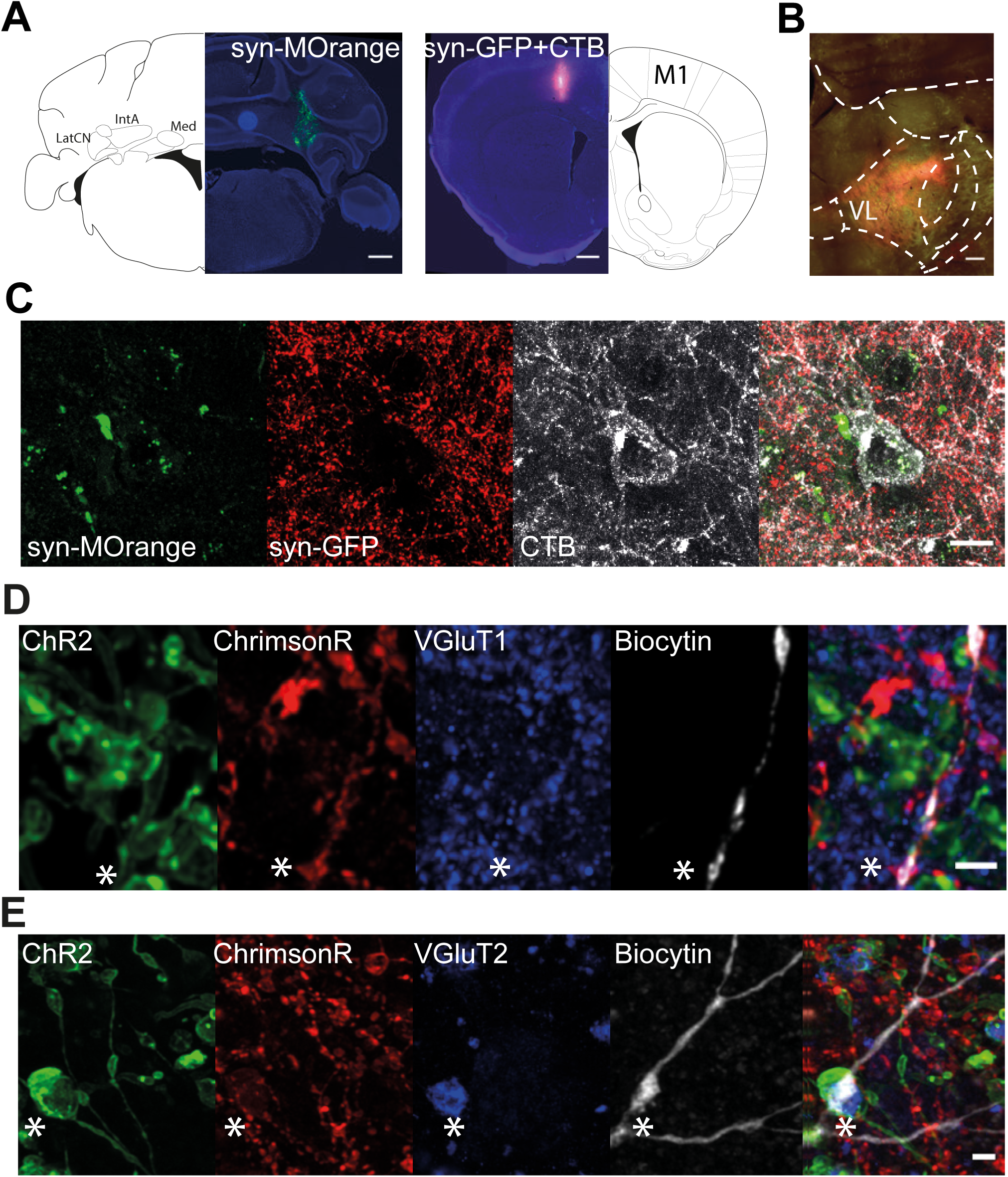
Morphological evaluation of the cerebello-thalamo-cortical connectivity. (A) Injection spots which represent Synaptophysin-mOrange expression in cerebellar nuclei (CN) and co-labelling of Synaptophysin-GFP and Cholera Toxin Subunit B (CTB) in primary motor cortex (M1) (scale bar: 500 µm). Synaptophysin-mOrange is represented in green and Synaptophysin-GFP in red. (B) The input from CN and M1 converges within VL nucleus and overlaps with retrogradely labeled (CTB positive) neurons that project back to M1 (scale bar: 500 µm). (C) Representative high-magnification image of a large volume cerebellar synapse and small volume M1 synapses that converge on a CTB-labeled VL neuron (white), which in turn projects back into the same area of M1 (scale bar: 10 µm). (D,E) ChrimsonR-positive fibers from M1-L6 and Channelrhodopsin-positive (ChR2) fibers from CN form synapses on the dendrites of biocytin-filled VL cells, which stain for vGluT1 or vGluT2, respectively. See Figure 3A for schematic representation of injection strategy. Terminals from M1-L6 (D) stain positive for vGluT1 and negative for vGluT2 (asterisks, scale bar: 2 µm), while CN synapses stain positive for vGluT2 (E) but not for vGluT1 (asterisks, scale bar: 2 µm). ChR2-YFP label CN fibers green, ChrimsonR-tdTomato labels M1-L6 fibers red, vGluT1 and 2 staining is depicted in blue and the biocytin-filled cell in white. Abbreviations: cerebellar nuclei (CN), lateral cerebellar nuclei (LatCN), anterior interposed nucleus (IntA), medial cerebellar nucleus (Med), primary motor cortex (M1), ventrolateral thalamic nucleus (VL).

To investigate the convergence of CN and M1-L6 inputs onto single VL cells electrophysiologically, we recorded their responses following dual optogenetic stimulation of AAV-ChR2-expressing CN fibers and AAV-flex-ChrimsonR-expressing M1 fibers from cortical layer 6 in *Ntsr1-Cre* mice (**Figure 3A**; (Klapoetke et al. 2014; Hooks et al. 2015). To ensure exclusive stimulation of M1-L6 or CN fibers in VL, we first evaluated photocurrents and spiking probabilities of ChrimsonR expressing M1-L6 neurons (**Supplementary Figure 3A-D**). After evaluating the impact of pulses at 585 nm (for 1, 15 and 200 ms) and 470 nm (for 1 ms), we settled on systematically recording reliable photocurrents evoked by 585 nm stimulations for 15 ms (**Supplementary Figure 3C-G**). In addition, we desensitized the ChrimsonR-expressing M1-L6 neurons to 470 nm photo stimulation by applying a pre-stimulation for 200 ms at 585 nm that saturates motor cortical photocurrents and prevents spike induction upon the subsequent 470 nm light pulse (**Supplementary Figure 3F,G;** Hooks et al., 2015; Klapoetke et al., 2014). Next, we showed that photo stimulation of ChR2 expressing CN terminals at 470 nm reliably evoked postsynaptic currents in VL neurons, whereas photo stimulation at 585 nm was insufficient to trigger synaptic release from these terminals (**Supplementary Figure 4A-D**). However, prolonged pre-stimulation at 585 nm before stimulation at 470 nm decreased cerebellar response amplitudes, when compared to stimulation with a single pulse at 470 nm (**Supplementary Figure 4F,G**). In summary, our *in-vitro* dual-optogenetic stimulation approach provides us with the unique opportunity to independently activate both pathways without inducing reverberating thalamo-cortical network oscillations such as found with *in-vivo* stimulation approaches.

**Figure 3.**
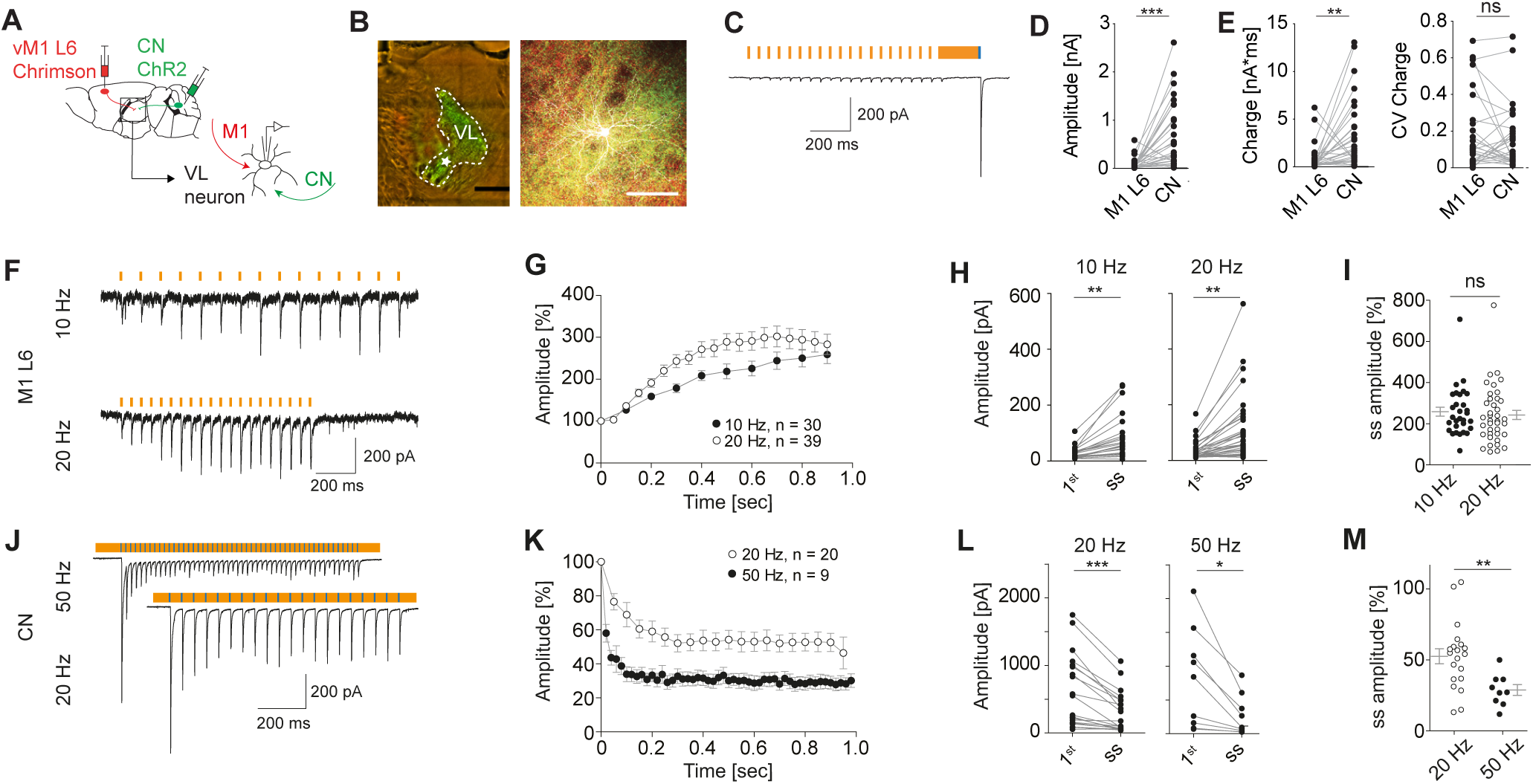
Physiological convergence of CN and M1 L6 in motor thalamus. (A) Schematic representation of the dual-optogenetic stimulation approach by expressing ChrimsonR-tdTomato (Chrimson) in motor cortical layer 6 (M1-L6) and ChannelRhodopsin-EYFP (ChR2) in cerebellar nuclei (CN) in Ntsr1-Cre transgenic mice. (B) Fluorescent images of Chrimson-positive M1-L6 fibers (red), ChR2-positive CN fibers (green) and a biocytin-filled VL neuron (white, scale bars: 100 µm). (C) Representative VL recording during 20 Hz M1-L6 stimulation (15 ms 585 nm, orange) and a single CN stimulus (200 ms 585 nm and 1 ms 470 nm, blue). The amplitude (D), charge and the coefficient of variation (CV) of the charge evoked by M1-L6 and CN stimuli (E). (F-H) Stimulus trains of 10 and 20 Hz (15 ms, 585 nm) selectively activate M1-L6 fibers and result in short-term facilitation in the steady state (ss) when compared to the first stimulus (1st). (I) The ss facilitation of M1-L6 inputs to VL neurons is not different between 10 and 20 Hz stimulus trains. (J-L) Stimulus trains of 20 and 50 Hz (1ms, 470 nm) in the presence of continuous 585nm light selectively activate CN fibers and result in short-term depression of VL EPSCs. (M) The ss depression of CN inputs to VL neurons is enhanced when comparing 50 to 20 Hz stimulus trains. ‘ns’ indicates not significant, ** P<0.01 and *** P<0.001.

We next evaluated the postsynaptic responses of individual cells in thalamic slices following activation of ChrimsonR-expressing M1-L6 fibers and ChR2-expressing CN fibers with light stimuli at 585 nm and 470 nm, respectively (**Figure 3B-E**). For M1-L6 stimulation the EPSC amplitudes were small at the beginning of the 20 Hz stimulation (18.6±4.0 pA), but they substantially increased towards the end of the steady state stimulation (89.7±24.5 pA). In contrast, stimulating CN fibers showed large initial EPSC amplitudes that decayed over time (688±131 pA, WCR-test CN vs. M1-L6 facilitated, *P*<0.0001, n=28; **Figure 3C-E**). The total charge transferred was 888±275 pA*ms for M1-L6 after a single stimulus in the facilitated state and 3308±708 pA*ms for initial responses from CN terminals (**Figure 3E**; WCR-test, *P*<0.0012, n=28), with a coefficient of variation (CV) of 0.19±0.03 and 0.16±0.03, respectively (**Figure 2E**; WCR-test, *P=*0.6406, n=28). Bath application of TTX decreased the post-synaptic responses evoked by M1-L6 and CN stimulation below noise level, in line with AP-driven neurotransmitter release (**Supplementary Figure 4H**). The postsynaptic response amplitudes after ChrimsonR or ChR2 activation were not correlated with incubation time after viral injection (**Supplementary Figure 4I**), indicating that the M1-L6- and CN-evoked EPSC amplitudes were independent of the expression level of the opsins. Moreover, all reconstructed dendritic trees resided within the anatomical borders of the VL (**Supplementary Figure 4J**, Paxinos and Franklin, 2001).

To further investigate the short-term synaptic response patterns of both inputs, we applied repetitive stimulation. After one second of 10 or 20 Hz stimulation M1-L6 EPSCs increased from 26.4±3.6 pA to 75.4±13.4 pA at 10 Hz and from 31.8±5.1 pA to 99.8±18.4 pA at 20 Hz (**Figure 3F-H**; WCR-test, *P*<0.0001 for 10 Hz (n=30) and 20 Hz (n=39)), corresponding to 258.8±21.7% and 242.8±22.2% of initial amplitudes, respectively. In contrast, train stimulation of CN fibers (20 or 50 Hz) resulted in decreased steady state EPSC amplitudes (**Figure 3J-L**, 302.5±67.7 pA at 20 Hz, WCR-test, *P*=0.0001, n=20; and 264.1±99.8 pA at 50 Hz, WCR-test, *P=*0.0039, n=9) corresponding to 52.5±5.3% and 28.8±3.8%, respectively. While the relative steady state amplitudes of M1-L6 evoked EPSCs showed no significant frequency modulation at 10 Hz and 20 Hz (**Figure 3G,I**; Mann-Whitney test, *P*=0.393, n=30 and 39, respectively), CN evoked EPSCs were significantly modulated by frequencies of 20Hz and 50Hz (**Figure 3K,M**; Mann-Whitney test, *P*=0.0039). These data provide the first direct evidence for the dichotomous excitatory inputs to individual VL neurons in which CN inputs evoke large initial but reduced steady state synaptic responses, whereas M1-L6 inputs evoke small but facilitating synaptic responses upon repetitive stimulation.

In the following step, we tested the interactions of cerebellar and cortical input patterns with thalamic membrane fluctuations, as well as the impact thereof on spiking behavior. In VL neurons innervated by both M1-L6 and CN inputs, M1-L6 inputs were insufficient to depolarize VL neurons above spiking threshold, whereas a single-pulse CN stimulus reliably induced spiking in 9 out of 11 cells (**Figure 4A**, 2.6±0.6 spikes per stimulus for CN). These data suggest that CN inputs drive the spiking response, while M1-L6 modulates the membrane potential. On average, stimulation of M1-L6 fibers at 10 Hz resulted in a steady state depolarization of 3.3±0.5 mV (**Figure 4B-D**; range 0.5-5.5 mV), which shifted the average membrane potential from −72.5±0.4 mV to −68.4±0.9 mV (**Figure 4E**; WCR-test, *P*=0.0039, n=9). Stimulation at 20Hz induced a depolarization of 3.5±0.5 mV (**Figure 4B-D**; ranging from 0.4 mV to 7.2 mV), resulting in a membrane potential shift from −72.8±0.6 mV to 69.1±0.8 mV (**Figure 4E**; WCR-test, *P*=0.0005, n=17). The depolarization induced by M1-L6 activation was independent of stimulation frequency (**Figure 4D**; Mann-Whitney test, *P*=0.7464).

**Figure 4.**
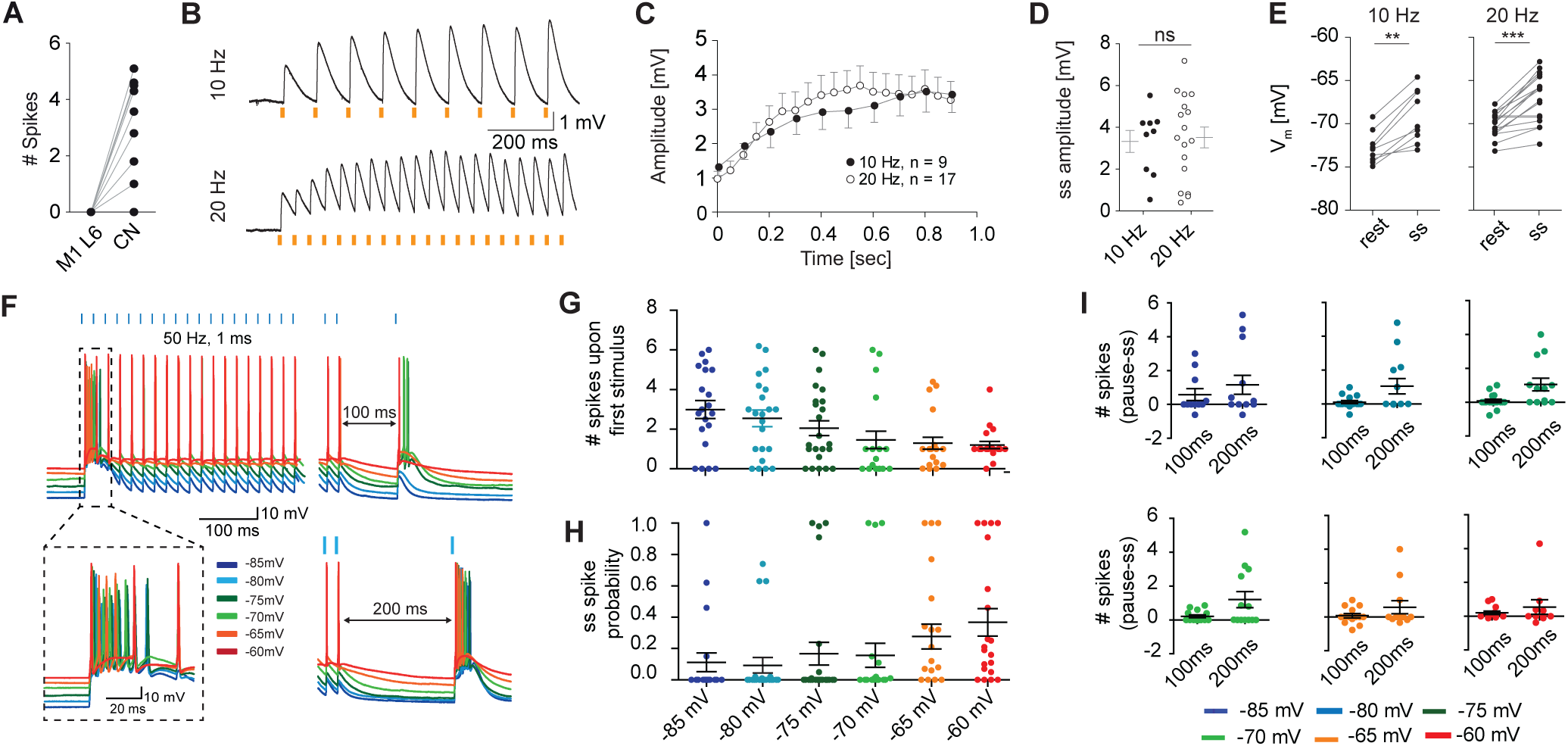
Motor cortex modulates VL membrane potential and cerebello-thalamic spike transfer. (A) CN stimulation (prolonged 585 nm and 1 ms 470 nm) induces spiking in most VL cells, while M1-L6 stimulation (15 ms, 585 nm) results in subthreshold depolarizations. (B) Example traces and (C) average membrane depolarizations evoked by 10 and 20 Hz M1-L6 stimulus trains (15 ms, 585 nm). (D,E) The steady state (ss) facilitation of M1-L6 inputs to VL neurons is not different between 10 and 20 Hz stimulus trains. (F) Example showing VL responses to 50 Hz CN stimulus trains (1 ms 470 nm, in absence of ChrimsonR-expression) and a subsequent 100 or 200 ms pause followed by a single CN stimulus. The increasing membrane potential reduces the number of action potentials fired upon the first CN-stimulus (G) and enhances the spike probability in the steady state (H). (I) Across membrane potentials, thalamic spike numbers increased after a pause of 100 and 200 ms compared to ss values. ** indicates P<0.01 and *** indicates P<0.001.

To decipher how the M1-L6 depolarization affects VL responses to CN-train stimulations and pauses, we injected depolarizing currents ranging within the steady-state amplitudes evoked by selective M1-L6 stimulation (11.3-562.6 pA and 0.4-7.2 mV), while stimulating ChR2 expressing CN fibers at 50 Hz (**Figure 4F**). With increasingly depolarized membrane potentials, the number of action potentials fired upon the first CN-stimulus decreased, but the spike probability following CN stimulation during the steady state increased (**Figure 4F-H**; number of spikes upon 1^st^ stimulus: 3.00±0.45 at −85 mV and 1.20±0.18 at −60 mV; steady state spike probability: 0.11±0.06 at −85 mV and 0.37±0.09 at −60 mV, Cochran-Armitrage test: *P*=0.0001 and 0.03267, respectively). Across membrane potentials, the number of spikes induced by stimulating CN fibers after pauses of 100 and 200 ms increased relative to the steady state spiking level (**Figure 4I**; 0.58±0.34 and 1.15±0.56 at −85 mV as well as 0.20±0.10 and 0.54±0.43 at −60 mV, respectively). Thus, a pause in high-frequency CN spiking activity is capable of re-inducing VL spiking, while the shift in membrane potential effectively modulates the gain of information transfer.

## Discussion

Our *in vitro* data reveal that attenuated responses to cerebellar stimulation in VL thalamic neurons are restored after a brief pause in the stimulus train. Moreover, we found that cerebral cortically mediated adaptation of the VL membrane potential determines not only the spiking probability during the stimulus train, but also the number of action potentials fired following pauses in CN stimulation. This synergistic modulation enables the motor thalamus to operate as a low-pass filter for the high-frequency cerebellar input that allows spike adaptation based on motor cortical feedback.

The high spontaneous firing frequencies of CN neurons, which can reach over 100 Hz in both *in vitro* and *in vivo* experimental settings (Raman et al. 2000; Hoebeek et al. 2010) are likely to deplete cerebellar synapses in the thalamus. Our results indeed show that optogenetic stimulation that synchronously activates ChR2-expressing CN axons up to 50 Hz results in short-term depression of EPSC amplitudes in all recorded VL neurons (**Supplementary Figure 2)**, which in turn dampens the transfer of information between VL and motor cortex. From this perspective, spike coding from thalamus to motor cortex is depending on recovery from synaptic depression during a pause in high-frequency CN spiking. Such pauses are found during several behavioral paradigms (Ohmae et al. 2013; Ten Brinke et al. 2017). For instance, during natural running on a treadmill the CN firing rate rises above 100 Hz during the lift of the paw and decreases sharply during the 100 ms before paw movement (Sarnaik and Raman 2018). According to our in-vitro findings, the transfer of cerebellar information from thalamus to cortex is attenuated during high-frequency CN activity, but can be relayed after the induction of initial spikes that follow a pause in CN spiking. Even though our in-vitro study can only assess the impact of synchronously activated cerebellar input with a limited maximum stimulus frequency (ChR2-stimulation of CN axons above 50 Hz resulted in failure to elicit neurotransmitter release; data not shown), the low-pass filter function within the thalamus suggests that cerebellar output is inversely encoded in VL. Whether the debated rebound spiking of CN neurons following a climbing fiber-mediated pause provides a significant contribution to thalamic information processing remains to be investigated (Gauck and Jaeger 2000; Alviña et al. 2008; Hoebeek et al. 2010; Bengtsson et al. 2011; Person and Raman 2011; Dykstra et al. 2016; Sarnaik and Raman 2018). Still, the properties that we describe in the current study already show that VL neurons translate the high-frequency cerebellar spiking pattern into a temporally sparse but precise and movement related code, which may serve to fine-tune the information processing in motor cortex.

The transfer of cerebellar information to the motor cortex is dictated by the state of the thalamic membrane potential (Jahnsen and Llinas 1984a; Mccormick and Bal 1997), which *in vivo* is determined by various subcortical modulatory inputs as well as intra- and extra-thalamic inhibitory inputs (Steriade et al. 1991; Giber et al. 2015; Halassa and Acsády 2016). Although our experimental setting does not account for such network effects, our findings show that by repetitive stimulation of M1 L6 fibers the thalamic membrane potential can be readily modulated and affect the availability of t-type calcium channels. Their activation drives thalamic cells to fire the characteristic low threshold calcium spike (LTS) and a burst of action potentials (Jahnsen and Llinas, 1984a, 1984b). The degree of t-type channel de-inactivation is time- and voltage-dependent and determines the number of spikes transferred within a burst (Jahnsen and Llinas 1984a; Mease et al. 2017) and thereby the quality and timing of thalamic spiking (Wolfart et al. 2005; Mease et al. 2014). It is important to note that in the in-vivo situation M1 L6 neurons mono-synaptically innervate excitatory neurons in VL as well as inhibitory neurons in the reticular thalamic nucleus (RTN), which in turn provide feed-forward inhibition to thalamo-cortical relay neurons (Yamawaki and Shepherd 2015). The depressing short-term release dynamics of RTN synapses in the thalamus shift the balance between excitation and inhibition induced by M1 L6 towards depolarized membrane potentials (Crandall et al. 2015). Future investigations should focus on unraveling the tri-partite integration of CN, M1 L6 and RTN inputs on VL neurons.

Our data show that the depolarizing shift in RMP after M1 L6 activation amplifies the cerebello-thalamic signal and enables the cortex to control the gain of thalamic processing in the motor system. As a consequence, the gain of spike transfer from VL to motor cortex does not only depend on the CN-evoked responses in VL neurons, but also on the synchronicity with M1 L6 synaptic background activity (Wolfart et al. 2005; Mease et al. 2014). The combination of cerebellar spike timing, response amplitude in VL neurons as well as their membrane potential can modulate the spike transfer to motor cortex along a continuum, as it has previously been shown for the sensory system in the awake mouse (Whitmire et al. 2016).

The motor cortical layer 5 neurons, which are essential for movement initiation and display changes in firing frequency correlative to movement execution (Brecht et al. 2004; Schiemann et al. 2015; Ebbesen et al. 2016; Sreenivasan et al. 2016), receive direct input from cerebellar-recipient VL neurons (Kuramoto et al. 2009; Hooks et al. 2013; Yamawaki and Shepherd 2015). Attenuating the transfer of cerebellar information to motor cortex by blocking spike transfer from VL to motor cortex results in a direct reduction in basal firing rate in layer 5 neurons, which eliminates movement-related adjustments in spiking-rate (Schiemann et al. 2015). The dependence of motor cortical activity on VL output highlights the essence of functional cerebello-thalamo-cortical signal integration for movement execution (Proville et al. 2014; Schiemann et al. 2015).

## Author contributions

C.B.S. performed all experimental work and analysis. C.B.S. and F.E.H. designed the experiments. F.E.H. and Z.G. provided technical support. F.E.H., Z.G. and C.I.D.Z. provided financial support. C.B.S. and F.E.H. wrote the original draft, Z.G. and C.I.D.Z. edited the manuscript. F.E.H. conceived and guided the project.

## Acknowledgements

We kindly thank Simona V. Gornati, Oscar H.J. Eelkman Rooda, Valentina Riguccini and Bas van Hoogstraten for fruitful scientific discussions and we thank Prof. Dr. Thomas Kuner for providing Synaptophysin virus constructs. We thank Erika H. Goedknegt, Elize D. Haasdijk, Mandy Rutteman, and Patrick Eikenboom for helping with immunohistochemical experiments. C.B.S. and F.E.H. are supported by the Dutch organization for life sciences (NWO-ALW; VIDI grant #016.121.346) and Medical Sciences (TOP-GO #91210067). Z.G. is supported by the Dutch organization for life sciences (NWO-ALM; VENI grant #863.14.001 and NWO-CAS grant #012.200.14) and the Erasmus MC fellowship. C.I.D.Z. thanks the Dutch Organization for Medical Sciences (Zon-MW; TOP-GO #91210067), Life Sciences (ALW; #854.10.004), ERC-adv (#294775) and ERC-POC (#768914) of the EU for support. None of the funding bodies had any input to the study design or outcome.

## Supplemetary Items

**Supplementary Figure 1:**

Representation and reconstruction of the individual injection locations and spread of viral transduction

**Supplementary Figure 2:**

Short-term dynamics and recovery after cerebellar stimulation

**Supplementary Figure 3:**

Photosensitivity of ChrimsonR-tdTomato expressing M1 L6 neurons

**Supplementary Figure 4:**

Independent excitation of CN fibers and properties of dually connected thalamic neurons

**Supplementary Table 1**:

Statistical analysis for all data

