## supplementary information for "Cerebello-Thalamic Spike Transfer via Temporal Coding and Cortical Adaptation"

Supplementary Figure 1

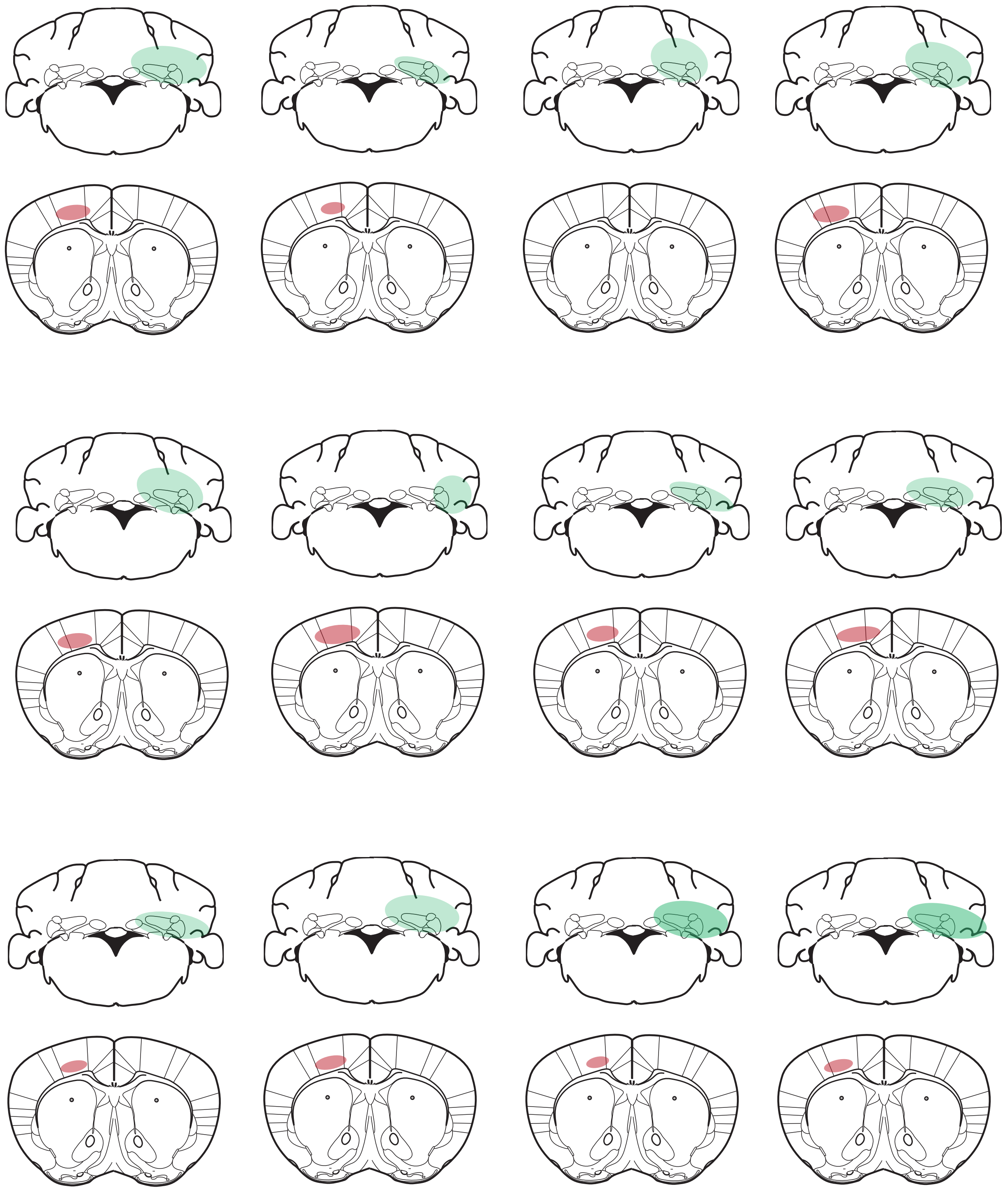

Supplementary Figure 2

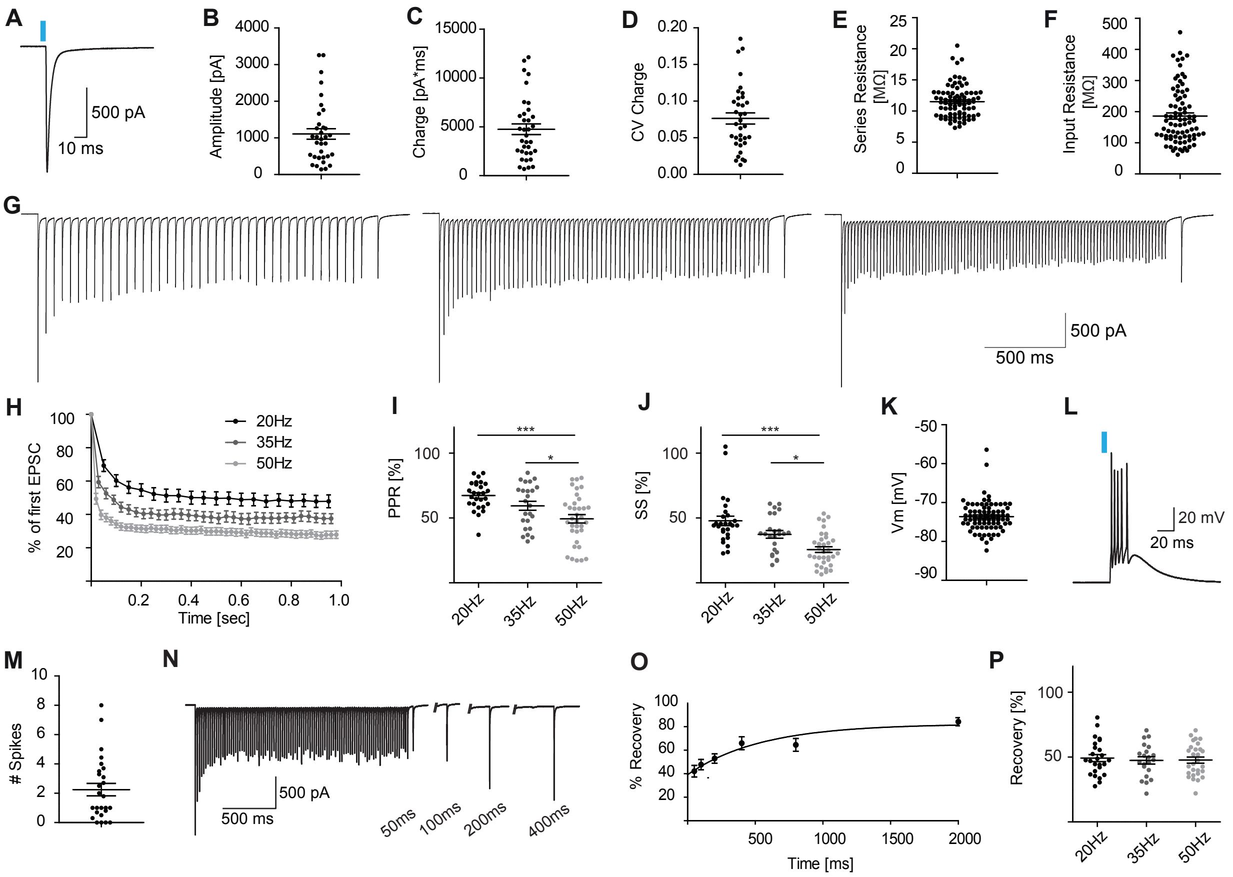

Supplementary Figure 3

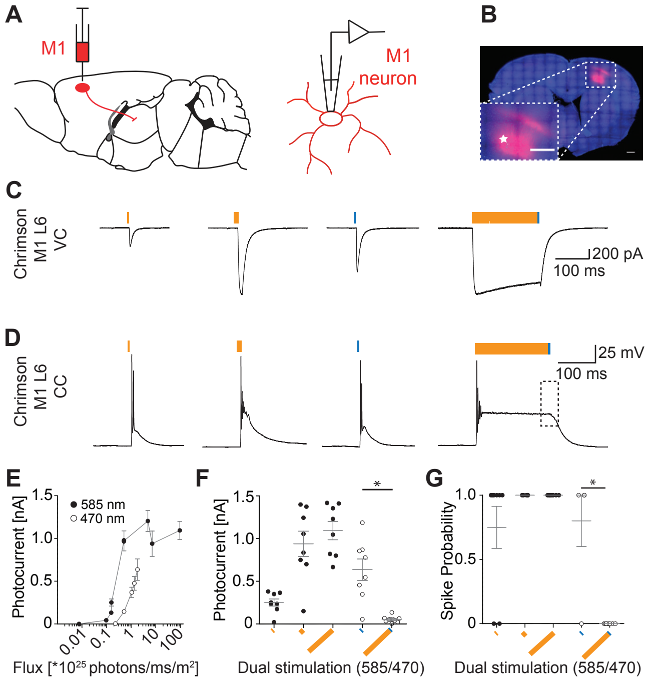

Supplementary Figure 4

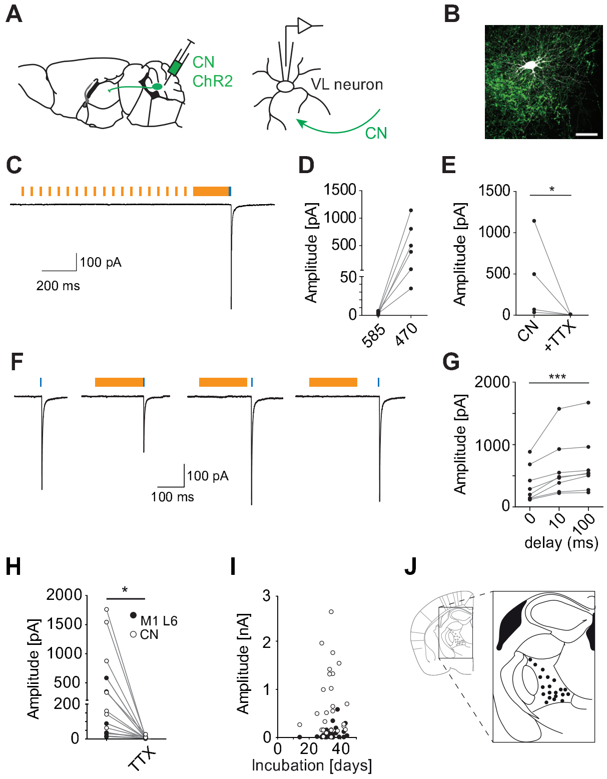

### Supplementary Figure Titles and Legends

**Supplementary Figure S1.** Representation and reconstruction of the individual injection locations and spread of viral transduction. ChR2-YFP was injected into the CN contralaterally to the recorded VL thalamus; ChrimsonR-tdTomato was injected into the M1 ipsilateral to the recorded VL.

**Supplementary Figure 2.** Short-term dynamics and recovery after cerebellar stimulation. (**A**) Example trace illustrating the optogenetically evoked response (1ms, 470nm) in a VL neuron. (**B-D**) The amplitude (**B**, 1107.0±143.6pA, n=35), charge (**C**, 4745.0±543.7pA*ms, n=35) and coefficient of variance (CV) of the charge (**D**, 0.08±0.01, n=35) for CN-evoked EPSCs are shown. The series resistance (**E**, 11.5±0.3MΩ, n=82) and input resistance (**F**, 186.2±10.0MΩ) are reported for all 82 recorded thalamic cells included in this study. (**G**) Example traces depicting CN-evoked responses to 20, 35 and 50Hz stimulus trains of 2s followed subsequently by an isolated 100ms pause and a single stimulus. (**H**) Normalized steady state (ss) depression evoked by 20, 35 and 50Hz stimulus train of 1s. (**I,L**) Repeated stimulation with 470nm at 20, 35 and 50Hz results in a paired-pulse ratio (PPR) of 67.3±2.1% (n=27), 59.8±3.4% (n=24) and 49.4±3.2% (n=35) (**I**, Friedman-test, 20Hz vs. 50Hz and 35Hz vs 50Hz *P*-values<0.001) and ss response of 47.8±3.6 (n=27), 37.4±2.9 (n=24) and 25.6±1.2% (n=35) (**L**, Friedman-test, 20Hz vs. 50Hz and 35Hz vs. 50Hz *P*-values<0.001). (**K**) The resting membrane potential ($V_m$) of VL neurons (-73.6±0.4mV, n=82). (**I**) The example trace depicts the thalamic burst response after cerebellar stimulation, which results in 2.24±0.42 spikes per stimulus (**M**, n=34). (**N**) The example trace depicts the time-dependent recovery of the compound cerebellar event, which is quantified in (**O**) (single exponential fit, R-square: 0.38, Plateau: 81.9%, Tau: 524.4ms). (**P**) The recovery of the CN-evoked response in VL neurons after 100ms pause was independent of frequency (20Hz: 49.2±2.7%, n=25; 35Hz: 47.4±2.8%, n=21; 50Hz: 47.7±2.4%, n=28; Friedman-test, *P*=0.324). * indicates *P*<0.05 and *** *P*<0.001.

**Supplementary Figure 3.** Photosensitivity of ChrimsonR-tdTomato expressing M1 L6 neurons. (**A**) Schematic illustration of ChrimsonR expression and patch-clamp recording from ChrimsonR expressing M1-L6 neurons. (**B**) Layer 6 specific expression of ChrimsonR-tdTomato in motor cortex of *Ntsr1-Cre* mice. (inset: high-magnification image of injection spot; scale bars 200µm). (**C,D**) Representative recordings of the photo-response from ChrimsonR-tdTomato expressing M1-L6 neurons after stimulation with 1, 15 and 200ms light at 585nm and 1ms at 470nm, when recorded in voltage-clamp (VC) (**C**) and current-clamp (CC) (**D**). To independently excite cerebellar synapses in thalamic slices (see Fig.2), the M1-L6 neuron was desensitized to light at 470nm by applying a 200ms stimulation at 585nm, following which a 1ms stimulus at 470nm failed to evoke an action potential in M1-L6 neurons (square in right panel of **D**). (**E,F**) Stimulating with pulses of 1, 15 and

200ms at 585 and 1ms at 470nm, which correspond to photonfluxes of $4.8*10^{24}$ (1ms) to $960*10^{24}$ (200ms) photons/ms/$m^2$ at 585nm and $18.08*10^{24}$ photons/ms/$m^2$ at 470nm (**E**) resulted in photocurrents that maximized in response to 15ms of 585nm (1ms 585nm: 251.1±42.1pA, 15ms 585nm: 938.8±148.3pA, 200ms 585nm: 1094±107pA, 1ms 470nm: 636.5±125.9pA, n=8 for all groups, **F**). The photocurrent in response to 1ms stimulation at 470nm is significantly decreased to 52.5±11.1pA after 200ms pre-stimulation at 585nm (**F**, Kruskal-Wallis test, $P<0.05$). (**G**) These maximal photocurrents induced AP firing in ChrimsonR positive M1-L6 neurons (1ms 585nm: 75±16%, n=8; 15ms 585nm: 100±0%, n=6; 200ms 585nm: 100±0%, n=8; 1ms 470nm: 80±20%, n=5), except when a 200ms 585nm pulse preceded the 1ms 470nm (0±0%, n=8, Kruskall-Wallis test, $P<0.05$). This dual-optogenetic stimulation paradigm prevents AP firing in M1-L6 neurons upon co-stimulation by 585nm and 470nm. * indicates $P<0.05$.

**Supplementary Figure 4.** Independent excitation of CN fibers and properties of dually connected thalamic neurons. (**A**) Schematic illustration of ChR2-expressing CN neurons and patch-clamp recording from VL neurons. (**B**) Fluorescent max-projection confocal image of a biocytin-filled VL neuron surrounded by ChR2-expressing CN fibers (scale bar: 50µm). (**C,D**) Example trace (**C**) and average post-synaptic response amplitude (**D**) evoked by a train of 585nm 15ms pulses and a 200ms block stimulation (average steady state EPSC amplitude after stimulation with 585nm: 3.7±0.7pA, n=6 and 1ms 470nm: 489.0±175.2pA, n=6), indicating that in our preparation 585nm stimulation does not trigger neurotransmitter release from ChR2-expressing CN fibers (Wilcoxon signed rank test, CN vs CN+TTX: $P=0.0313$) (**E**) The cerebellar release is AP-dependent as the response is fully blocked by application of 10µM TTX (CN: 436.0±258.0pA, with TTX: 6.6±2.3pA, n=4). (**F,G**) To control for the interaction between stimulation at 585nm and 470nm, we evaluated the effect of 585nm stimulation on the responses evoked by 470nm stimulation. Delaying the 470nm pulse by 10ms after cessation of the 585nm pulse increased the EPSC amplitude significantly (0ms delay: 360.0±101.1pA, 10ms delay: 607.4±158.7pA, 100ms delay: 663.3±164.9pA, Friedman test, 0ms delay vs 100ms delay: $P<0.0001$, n=8). (**H**) The release from both CN (696.2±267.7pA) and M1-L6 (160.3±83.9pA) is blocked after application of tetrodotoxin (TTX; 9.1±3.2pA and 5.1±1.5pA, respectively, n=7, $P=0.015$, Wilcoxon signed rank test). (**I**) The postsynaptic response amplitude evoked by ChrimsonR or ChR2 activation was not correlated with incubation time (linear regression, M1-L6: $r^2=0.1003$, CN: $r^2=0.0045$, n=28). (**J**) The location of most reconstructed neurons (20 out of 28 dually connected neurons) was mapped and projected to a single plane (1.2mm posterior of Bregma) of VL in the Paxinos Atlas. * indicates $P<0.05$ *** indicates $P<0.001$

| **Supplementary Table 1** Statistical analysis for all data in Figure 1 | | | | | | | |
|---|---|---|---|---|---|---|---|
| **Panel** | **Unit** | **Mean ± SEM** | **Normal distribution** | **Pairing** | **Test applied** | **P-value** | **N** |
| 1c | Spike probability | 20Hz: 0.20 ± 0.08<br>50Hz: 0.15 ± 0.07 | No<br>No | Yes<br>Yes | Wilcoxon signed rank test | 0.0625 | 25<br>25 |
| 1d | # spikes at resting potential | 50ms: 0.15 ± 0.09<br>100ms: 0.28 ±0.1<br>200ms: 1.39 ± 0.46<br>400ms: 2.43 ± 0.65 | No<br>No<br>No<br>Yes | Yes<br>Yes<br>Yes<br>Yes | Friedman test, Bonferroni correction for multiple comparisons | 50ms vs. 100ms =1.0<br>100ms vs. 200ms =1.0<br>50ms vs. 200ms =1.0<br>50ms vs. 400ms =0.171<br>200ms vs. 400ms =0.602<br>100ms vs. 400ms =0.019 | 15<br>19<br>18<br>14 |

| **Supplementary Table 2** Statistical analysis for all data in Figure 3 | | | | | | | |
|---|---|---|---|---|---|---|---|
| **Panel** | **Unit** | **Mean ± SEM** | **Normal distribution** | **Pairing** | **Test applied** | **P-value** | **N** |
| 3d | Amplitude [pA] | CN: 688.4 ± 130.7<br>M1 L6: 89.7 ± 24.5 | No<br>No | Yes | Wilcoxon signed rank test | <0.0001 | 28 |
| 3e | Charge [pA*ms] | CN: 3308.0 ± 708.0<br>M1 L6: 888.5 ± 275.3 | No<br>No | Yes | Wilcoxon signed rank test | 0.0012 | 28 |
| 3e | CV | CN: 0.16 ± 0.03<br>M1 L6: 0.19 ± 0.04 | No<br>No | Yes | Wilcoxon signed rank test | 0.6406 | 28 |
| 3h | Amplitude [pA] | 10Hz $1^{st}$: 26.4 ± 3.6<br>10Hz Ss: 75.4 ± 13.4 | No<br>No | Yes | Wilcoxon signed rank test | <0.0001 | 30 |
| | | 20Hz $1^{st}$: 31.9 ± 5.1<br>20Hz Ss: 99.8 ± 18.4 | No<br>No | Yes | Wilcoxon signed rank test | <0.0001 | 39 |
| 3i | Amplitude [%] | 10Hz: 258.8 ± 21.7<br>20Hz: 242.8 ± 22.2 | No<br>No | No | Mann Whitney test | 0.3934 | 30<br>39 |
| 3l | Amplitude [pA] | 20Hz $1^{st}$: 603.1 ± 119.4<br>20Hz Ss: 302.5 ± 67.7 | Yes<br>No | Yes | Wilcoxon signed rank test | 0.0001 | 20 |
| | | 50Hz: $1^{st}$: 811.1 ± 242.8<br>50Hz Ss: 264.1 ± 99.8 | Yes<br>Yes | Yes | Wilcoxon signed rank test | 0.0039 | 9 |
| 3m | Amplitude [%] | 20Hz: 52.5 ± 5.3<br>50Hz: 28.8 ± 3.8 | Yes<br>Yes | No | Mann Whitney test | 0.0032 | 20<br>9 |

| **Supplementary Table 3** Statistical analysis for all data in Figure 4 | | | | | | | |
|---|---|---|---|---|---|---|---|
| **Panel** | **Unit** | **Mean ± SEM** | **Normal distribution** | **Pairing** | **Test applied** | **P-value** | **N** |
| 4a | # Spikes | CN: 2.6 ± 0.6<br><br>M1 L6: 0 | Yes | NA | NA | NA | 11 |
| 4d | Amplitude [mV] | 10Hz: 3.3 ± 0.5<br>20Hz: 3.5 ± 0.5 | Yes<br>Yes | No | Mann Whitney test | 0.7464 | 9<br>17 |
| 4e | Vm [mV] | 10Hz rest: -72.8 ± 0.6<br>10Hz Steady state: -69.1 ± 1.0 | Yes<br>Yes | Yes | Wilcoxon signed rank test | 0.0039 | 9 |
| | | 20Hz rest: -72.5 ± 0.4<br><br>20Hz Steady state: -68.4 ± 0.86 | Yes<br><br>Yes | Yes | Wilcoxon signed rank test | 0.0005 | 17 |
| 4g | # spikes | -85mV: 3.00 ± 0.45<br>-80mV: 2.55 ± 0.42<br>-75mV: 2.01 ± 0.37<br>-70mV: 1.64 ± 0.45<br>-65mV: 1.29 ± 0.30<br>-60mV: 1.20 ± 0.18 | Yes<br>Yes<br>No<br>No<br>No<br>No | No<br>No<br>No<br>No<br>No<br>No | Cochran Armitrage Trend Test | 0.0001 | 20<br>22<br>25<br>21<br>21<br>20 |
| 4h | Spike probability | -85mV: 0.11 ± 0.06<br>-80mV: 0.09 ± 0.05<br>-75mV: 0.17 ± 0.07<br>-70mV: 0.16 ± 0.08<br>-65mV: 0.28 ± 0.08<br>-60mV: 0.37 ± 0.09 | No<br>No<br>No<br>No<br>No<br>No | No<br>No<br>No<br>No<br>No<br>No | Cochran Armitrage Trend Test | 0.03267 | 20<br>22<br>25<br>21<br>21<br>20 |
| 4i | # spikes (pause-ss) | -85mV 100ms: 0.58 ± 0.34<br>-85mV 200ms: 1.15 ± 0.56<br>-80mV 100ms: 0.10 ± 0.09<br>-80mV 200ms: 1.05 ± 0.45<br>-75mV 100ms: 0.07 ± 0.10<br>-75mV 200ms: 1.04 ± 0.38<br>-70mV 100ms: 0.21 ± 0.09<br>-70mV 200ms: 1.20 ± 0.48<br>-65mV 100ms: 0.08 ± 0.13<br>-65mV 200ms: 0.58 ± 0.38<br>-60mV 100ms: 0.20 ± 0.10<br>-60mV 200ms: 0.54 ± 0.43 | No<br>No<br>No<br>No<br>No<br>No<br>No<br>No<br>No<br>No<br>No<br>No | No<br>No<br>No<br>No<br>No<br>No<br>No<br>No<br>No<br>No<br>No<br>No | | | 11<br>13<br>16<br>13<br>17<br>13<br>14<br>13<br>14<br>12<br>13<br>10 |

**Supplementary Table 4** Statistical analysis for all data in Supplementary Figure 2

| **Panel** | **Unit** | **Mean ± SEM** | **Normal distribution** | **Pairing** | **Test applied** | **P-value** | **N** |
|---|---|---|---|---|---|---|---|
| 2b | Amplitude [pA] | 1107.0 ± 143.6 | No | NA | NA | NA | 35 |
| 2c | Charge [pa*ms] | 4745.0 ± 543.7 | Yes | NA | NA | NA | 35 |
| 2d | CV | 0.076 ± 0.007 | Yes | NA | NA | NA | 35 |
| 2e | Series Resistance [MΩ} | 11.5 ± 0.3 | No | NA | NA | NA | 82 |
| 2f | Input Resistance [MΩ] | 186.2 ± 10.0 | No | NA | NA | NA | 82 |
| 2i | PPR [%] | 20Hz: 67.3 ± 2.1<br>35Hz: 59.3 ± 3.4<br>50Hz: 49.4 ± 3.2 | Yes<br>Yes<br>Yes | No<br>No<br>No | Friedman test, Bonferroni correction for multiple comparisons | 20Hz vs 35Hz =0.789<br>20Hz vs 50Hz <0.001<br>35Hz vs 50Hz <0.001 | 27<br>24<br>35 |
| 2j | SS [%] | 20Hz: 47.8 ± 3.6<br>35Hz: 37.8 ± 2.9<br>50Hz: 25.6 ± 2.2 | No<br>Yes<br>Yes | No<br>No<br>No | Friedman test, Bonferroni correction for multiple comparisons | 20Hz vs 35Hz =0.262<br>20Hz vs 50Hz <0.001<br>35Hz vs 50Hz <0.001 | 27<br>24<br>35 |
| 2k | Resting Potential [mV] | -73.6 ± 0.4 | No | NA | NA | NA | 82 |
| 2m | # spikes | 2.24 ± 0.42 | No | NA | NA | Na | 26 |
| 2p | Percent recovery | 50ms: 42.1 ± 4.9<br>100ms: 47.7 ± 4.4<br>200ms: 52.9 ± 4.0<br>400ms: 65.9 ± 5.5<br>800ms: 64.5 ± 5.5<br>2000ms: 84.2 ± 3.5 | Yes<br>Yes<br>Yes<br>Yes<br>Yes<br>Yes | No<br>No<br>No<br>No<br>No<br>No | Single Exponential fit | Y0: 39.7<br>Plateau: 81.9<br>Tau: 524.4<br>$R^2$: 0.38 | 12<br>11<br>12<br>12<br>6<br>5 |
| 2o | Recovery [%] | 20Hz: 49.2 ± 2.7<br>35Hz: 47.4 ± 2.8<br>50Hz: 47.7 ± 2.4 | Yes<br>Yes<br>Yes | No<br>No<br>No | Friedman test, Bonferroni correction for multiple comparisons | 0.324 | 25<br>21<br>28 |

**Supplementary Table 5** Statistical analysis of all data in Supplementary Figure 3

| **Panel** | **Unit** | **Mean ± SEM** | **Normal distribution** | **Test applied** | **P-value** | **N** |
|---|---|---|---|---|---|---|
| 3f | Photo-current [nA] | 1ms 585nm:<br>251.5 ± 42.1<br>15ms 585nm:<br>938.8 ± 148.3<br>200ms 585nm:<br>1094 ± 106.8<br>1ms 470nm:<br>636.5 ± 125.9<br>585nm+470nm:<br>52.5 ± 11.1 | Yes<br><br>Yes<br><br>Yes<br><br>Yes<br><br>No | Friedman test, Bonferroni correction for multiple comparisons | 1ms 585nm vs. 15ms 585nm:<br>=0.339<br>1ms 585nm vs. 200ms 585nm:<br>=0.133<br>1ms 585nm vs. 1ms 470nm:<br>=1.0<br>1ms 585nm vs. 585nm + 470nm:<br>=0.477<br>15ms 585nm vs. 200ms 585nm:<br>=1.0<br>15ms 585nm vs. 1ms 470nm:<br>=1.0<br>15ms 585nm vs. 585nm + 470nm:<br>< 0.001<br>200ms 585nm vs. 1ms 470nm:<br>=1.0<br>200ms 585nm vs. 585nm + 470nm<br>< 0.001<br>1ms 470nm vs. 585nm + 470nm:<br>=0.003 | *8*<br><br>*8*<br><br>*8*<br><br>*8*<br><br>*10* |
| 3g | Spike probability | 1ms 585nm:<br>0.75 ± 0.16<br>15ms 585nm:<br>1.0 ± 0<br>200ms 585nm:<br>1.0 ± 0<br>1ms 470nm:<br>0.8 ± 0.2<br>585nm+470nm:<br>0 ± 0 | Yes<br><br>NA<br><br>NA<br><br>NA<br><br>NA | Friedman test, Bonferroni correction for multiple comparisons | 1ms 585nm vs. 15ms 585nm:<br>=1.0<br>1ms 585nm vs. 200ms 585nm:<br>=1.0<br>1ms 585nm vs. 1ms 470nm:<br>=1.0<br>1ms 585nm vs. 585nm + 470nm:<br>=0.328<br>15ms 585nm vs. 200ms 585nm:<br>=1.0<br>15ms 585nm vs. 1ms 470nm:<br>=1.0<br>15ms 585nm vs. 585nm + 470nm:<br>=0.09<br>200ms 585nm vs. 1ms 470nm:<br>=1.0<br>200ms 585nm vs. 585nm + 470nm:<br>=0.057<br>1ms 470nm vs. 585nm + 470nm:<br>=0.012 | *8*<br><br>*6*<br><br>*8*<br><br>*5*<br><br>*8* |

| Supplementary Table 6 Statistical analysis of all data in Supplementary Figure 4 | | | | | | | |
|---|---|---|---|---|---|---|---|
| **Panel** | **Unit** | **Mean ± SEM** | **Normal distribution** | **Pairing** | **Test applied** | **P-value** | **N** |
| 4d | Amplitude [pA] | 585nm:<br>3.7 ± 0.8<br>470nm:<br>489.0 ± 175.2 | Yes<br><br>Yes | NA<br><br>NA | Wilcoxon signed rank test | 0.0313 | 6<br><br>6 |
| 4e | Amplitude [pA] | CN:<br>436.0 ± 258.1<br>CN + TTX:<br>6.5 ± 2.3 | NA<br><br>NA | NA<br><br>NA | NA | NA<br><br>NA | 4<br><br>4 |
| 4g | Amplitude [pA] | 0ms:<br>360 ± 101.1<br>10ms:<br>607.4 ± 158.7<br>100ms:<br>663.3 ± 164.9 | Yes<br><br>No<br><br>No | NA<br><br>NA<br><br>NA | Friedman Test And Dunn's Multiple Comparisons Test | 0ms vs. 10ms:<br>>0.05<br>10ms vs. 100ms<br>>0.05<br>0ms vs. 100ms:<br><0.0001 | 8<br><br>8<br><br>8 |
| 4i | Amplitude [pA] | M1 L6: 160.3 ± 83.8<br>M1 L6 TTX: 5.1 ± 1.5<br>CN: 696.2 ± 267.7<br>CN TTX: 9.1 ± 3.2 | NA<br>NA<br>NA<br>NA | Yes<br>Yes<br>Yes<br>Yes | Wilcoxon signed rank test | 0.0156<br><br>0.0156 | 7<br>7<br>7<br>7 |
| 4j | Amplitude vs Incubation | M1 L6<br>CN | NA<br>NA | Yes<br>Yes | Linear regression | $R^2$:<br>0.1003<br>0.0045 | 28<br>28 |
